# Individual Alpha Frequency Predicts the Sensitivity of Time Perception

**DOI:** 10.1101/2024.12.16.628734

**Authors:** Audrey Morrow, Montana Wilson, Michaela Geller-Montague, Sara Soldano, Sabah Hajidamji, Jason Samaha

**Affiliations:** University of California, Santa Cruz, Santa Cruz, CA 95064; University of California, San Francisco, San Francisco, CA 94143; Stanford University, Stanford, CA 94305

**Keywords:** alpha oscillations, individual differences, duration estimation, duration discrimination, time perception

## Abstract

A growing body of research links individual differences in the alpha-band frequency to temporal aspects of perception. However, whether the human alpha rhythm is a correlate of time perception itself has remained controversial. This study combined EEG with multiple duration perception tasks to evaluate whether individual alpha frequency (IAF) is associated with sensitivity or bias in judging visual durations across a range of peri-second durations (spanning 100-1200ms). In a temporal estimation task, participants (n = 55; 38 female, 13 male, 4 non-binary) reported the duration of a single stimulus between 300-1200ms. In a temporal discrimination task, participants reported which of two stimuli was longer: a standard (100, 600, or 1200ms) or comparison (50-150% of the standard). Stimuli also varied in whether their luminance was static or dynamically varying over time. We found that IAF was significantly related to the variance of duration estimates, a precision measure, but not average duration estimates, a bias measure. Further supporting this relationship, psychometric function slopes obtained from the independent duration discrimination tasks were positively correlated with IAF, particularly for the static stimulus conditions. These individual differences effects held when controlling for participant age. We also explored trial-level variability in alpha frequency and found it was predictive of shifts in the point of subjective equality (PSE) during discrimination of very short (100ms). Taken together, these results suggest that IAF plays a role in shaping individual differences in the sensitivity of time perception and that spontaneous variations around one’s IAF can lead to a bias in temporal representations.

**Significance Statement:** Brain waves in the 8-13 Hz range, known as alpha waves, have long been hypothesized to modulate our perception of time, yet the evidence remains unclear. This study investigates the relationship between an individual’s alpha frequency (IAF) and temporal sensitivity using a wide range of time perception tasks and stimulus durations to address gaps in the literature. We demonstrate that IAF is significantly associated with the precision of duration estimates and sensitivity in duration discrimination, particularly for static unchanging stimuli. These findings provide novel evidence that IAF shapes individual differences in time perception, emphasizing its role as a neural marker of temporal sensitivity.

## Introduction

The neural mechanisms underlying the perception of stimulus duration have long intrigued psychologists since, unlike other sensory features, time has no dedicated sensing organ. One view is that intrinsic neural dynamics act as an internal clocking mechanism that contributes to the perceived passage of time (Ivry & Schlerf, 2008). Alpha-band (8-13 Hz) neural oscillations have garnered interest as a potential neural clocking mechanism given their relatively stable rhythmic characteristics, intrinsic genesis, and the fact that individual variation in alpha frequency has been associated with a range of visual and cross-modal temporal integration processes in perception (Cecere et al., 2015; Dou et al., 2022; Drewes et al., 2022a; Keil & Senkowski, 2017; Migliorati et al., 2020; Samaha & Postle, 2015; Samaha & Romei, 2024; Venskus & Hughes, 2021). Indeed, early researchers debated the idea that the alpha rhythm reflects the minimum “psychological moment” (Ellingson, 1956; Treisman, 1984; White, 1963). If true, the frequency of occurrence of these moments, as indexed by an individual’s alpha frequency (IAF) may predict individual variation in duration perception ability.

Early research on the link between alpha frequency and time perception (reviewed in van Wassenhove et al., 2019) led to mixed results, with several reports of small-to-medium correlations between IAF and time estimates (Cahoon, 1969; Werboff, 1962) but also null findings (Legg, 1968). However, a major limitation of these earlier studies is that the measures of time perception that were used often did not distinguish between bias (over/under estimation of time) and sensitivity/precision (the ability to reliably distinguish intervals). And, early work often used durations spanning several seconds up to minutes, which arguably reflect more mnemonic, rather than perceptual, contributions to performance.

More recent work has overcome these limitations by examining the connection between IAF and bias and sensitivity using shorter, peri-second durations, but has also led to inconsistent findings. A study by Milton and Pleydell-Pearce (2016) used a two-interval forced choice (2IFC) temporal discrimination task to explore links between IAF and prestimulus alpha phase on bias and sensitivity. They found that IAF and phase were related to bias in perception of durations around 400ms, as measured by the point of subjective equality (PSE), but IAF was not related to sensitivity, as measured by the slope of individual psychometric functions. Another recent study by Mioni et al. (2020) used tACS at 2Hz +/- IAF during a temporal generalization task and found that stimulation above (below) IAF produced a bias towards longer (shorter) judgments of visual stimuli compared to a 500ms memorized standard. However, they did not find an effect of stimulation on the precision of perceptual judgments. In contrast to these results on bias, Mokhtarinejad et al. (2024) used a very similar temporal generalization task to Mioni et al. (2020) except with a 1000ms standard and found a positive correlation between IAF and precision, but not bias. However, they also found no effect of tACS stimulation above or below IAF. One possible source of the variability in these findings is the use of relatively small sample sizes for individual differences research. Indeed, the three studies reviewed above used samples between 15 and 24 participants, leaving open the possibility that discrepant results reflect sampling error. Thus, research to date has yet to measure IAF and peri-second time perception bias and sensitivity in a relatively larger sample of participants.

Here we derived measures of both sensitivity (precision) and bias in peri-second temporal *estimation*, which requires judging stimulus durations relative to an internal reference, and in temporal *discrimination*, which requires comparative judgment of two external stimulus durations, in a sample of 55 participants. We identified IAF from eyes-closed resting-state EEG recordings and observed positive correlations between IAF and precision in the duration estimation task and sensitivity in the discrimination task, but no relationship with bias at the individual level. Within-subjects analyses showed a small but significant correlation between instantaneous alpha frequency and PSE in duration discrimination. Our findings suggest that individuals with higher alpha frequencies have more sensitive time perception, but fluctuations around IAF may lead to a bias.

## Method

### Participants

This study was approved by the University of California Santa Cruz (UCSC) Institutional Review Board. Fifty-five participants with a mean age of 21.46 (SD = 5.16, range: 18-35; 38 female, 13 male, 4 non-binary) were recruited from UCSC’s online participant portal (SONA Systems) and from the Samaha Lab. This sample size was chosen to achieve 90% power to detect a correlation in the range suggested by a recent meta-analysis of the correlation between IAF and temporal binding measures (mean r ∼ 0.45; Samaha & Romei, 2024). The study consisted of two days of testing; the first day of testing took no more than 1.5 hours and involved the completion of questionnaires and behavioral tasks. Participants received 1.5 research credits and a $10 Amazon gift card. The second day of testing, which had to be completed within 12 days of the first, took no more than 3.5 hours and involved the completion of several duration perception tasks with simultaneous EEG recording. Participants were rewarded with 3.5 research credits and a $30 Amazon gift card. The sample consisted of 68.52% of participants who identified as female, 24.07% who identified as male, and 7.41% who identified as non-binary. The participant sample was 45.45% White/Caucasian, 25.45% Asian (35.71% did not specify further, but of those who did specify, Indian-identifying participants made up 14.29% and those who identified as either Chinese, Cambodian, Filipino, Japanese, Persian, Taiwanese, or Vietnamese each made up 7.14%), 20% Hispanic/Latino, and 9.09% multiracial (20% Black/Southeast Asian, 20% White/Indian, 20% White/Filipino, 20% White/Taiwanese, and 20% White/Chinese). Participants all reported having normal or corrected-to-normal vision.

#### Duration Estimation Task

In the duration estimation task participants provided estimates of the duration of a simple visual stimulus (Figure 1A). In both the practice and experimental blocks, participants observed a dark gray dot presented on a medium gray background at either 3 degrees of visual angle (DVA) above or below the central fixation. The dot duration was pseudo-randomly chosen between 300-1100ms in 100ms intervals with equal probability. Participants estimated each stimulus duration using a slider, controlled via computer mouse, that displayed a number line of possible durations, ranging from 100-1400ms in 100ms intervals. The slider provided numbers outside of the range of actual durations to reduce bias when reporting the shortest and longest possible durations. The current value was displayed on the screen in numerical values and was represented on the slider by a green dot which appeared at a random starting location on each trial. The inter-trial interval (ITI) ranged between 1.2-1.8s after the response was input. The practice block was 48 trials long and provided feedback displaying the actual stimulus duration if participants were incorrect in their estimate or “Correct!” if participants were correct. The two experimental blocks lacked feedback and were 90 trials long with each possible duration presented 10 times, totaling 180 trials.

**Figure 1:**
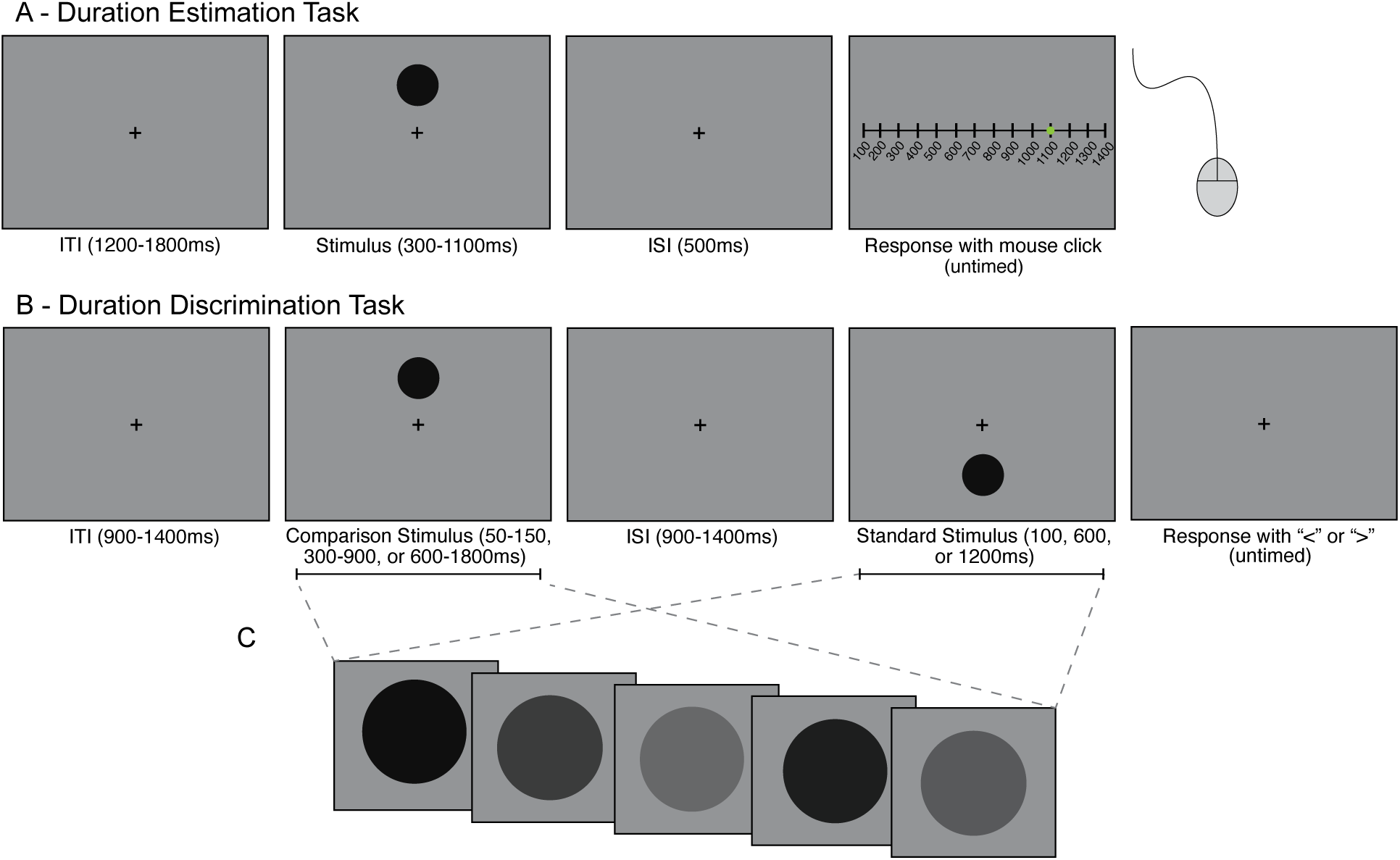
Diagram of duration perception tasks. (A) Duration estimation task. Participants were shown a dot of variable duration either above or below the central fixation and reported their estimate of the duration of the dot via mouse click along a slider of possible durations. (B) Duration discrimination task. Participants were presented with a dot above fixation, followed by a dot below fixation, or vice versa (dot presentation alternated by block) and reported, via button press, whether the first or second dot was longer. (C) A schematic of the dynamic stimulus condition. The luminance value of the stimulus in the dynamic version of the task varied randomly between black and gray on every screen refresh (120 Hz) throughout the stimulus presentation.

#### Duration Discrimination Task

The duration discrimination task was a two-interval forced choice (2IFC) design in which participants were asked to report which of two visual stimuli they perceived to be longer in duration (Figure 1B-C). This task had two conditions: a “static” condition, where the stimulus was the same gray dot as presented in the duration estimation task, and a novel “dynamic” condition where the dot varied randomly in luminance between medium gray (0.5 luminance) to black (0 luminance) on every monitor refresh (120 Hz). There is evidence that changes in luminance are cross correlated with the broadband EEG signal in a manner related to the participant’s IAF (VanRullen & Macdonald, 2012), suggesting that a dynamically modulating stimulus should elicit a strong response at the alpha frequency and potentially strengthen any link between IAF duration discrimination. To make the dynamically modulating stimulus, we created random luminance sequences for each dot presentation on each trial, and then normalized the sequence amplitude in the Fourier domain before performing an inverse Fourier transform to ensure the dot had equal energies at all frequencies. Besides the introduction of a flickering dot, the dynamic task was identical to the static task. The order of these conditions was counterbalanced across experimental sessions.

In both duration discrimination tasks, participants observed the first dot either 3 DVA above or below the central fixation, and then a second target on the opposite side of fixation (to avoid any adaptation effects), with the presentation order counterbalanced across blocks so that participants could always expect the location of the first dot as well as the second dot. In other words, whether the first stimulus was presented above or below fixation alternated from block to block but was held constant across the block. For all blocks, one target (the “standard”) was presented for a standard duration of 100, 600, or 1200ms, and the other target (the “comparison”) was presented for a fraction of the standard duration (either 50%, 70%, 90%, 110%, 130%, or 150% of the standard duration). In this way, the comparison was always proportional to the standard, scaling the comparison durations to be longer for the longer standard durations and shorter for the shorter standards in order to equate difficulty for the different standard durations as predicted by Weber’s Law (Haigh et al., 2021). There was a total of 30 presentations of each standard and comparison pair for each task condition. After participants responded with a button press to indicate whether the first (“<”) or the second (“>”) target was longer in duration, there was an inter-trial interval of 0.9-1.4s. The practice blocks used comparison durations that were 50%, 80%, 120%, and 150% to give participants exposure to some of the easier discriminations and allow them to learn the task, as well as some discriminations of medium difficulty as practice for the experimental block. There were 48 practice trials total where participants received feedback in the form of a tone if they selected the wrong stimulus. The main task comprised 540 trials for both the static and dynamic versions of the task. For all blocks, the presentation order of the standard and comparison varied each trial.

#### Critical Flicker Frequency (CFF)

The CFF - defined as the frequency at which continuously flickering illumination is perceived as constant - was used as a marker of an individual’s temporal resolution in their vision (Cass et al., 2011; Eisen-Enosh et al., 2017). Temporal resolution is an aspect of visual processing that is often correlated with IAF in seemingly related tasks such as the two-flash fusion illusion (Drewes et al., 2022b; Samaha & Postle, 2015). Individual differences in CFF are also related to IAF in clinical populations with hepatic encephalopathy (Baumgarten et al., 2018; Butz et al., 2013; May et al., 2014). We thus explored whether the CFF task might relate to IAF as well as the other duration perception measures in our non-clinical sample, and administered this task across both days of testing. The CFF was measured with the Flicker-Fusion system by Lafayette Instruments, which is designed to provide a quick (1-2 minutes) measure of an individual’s CFF. Participants viewed a flashing light through a viewing chamber and reported whether they perceived the light as flashing or continuous. The light was presented binocularly at the same rate to each eye and the rate of presentation adaptively changed according to participant responses. The initial flicker frequency was 20Hz, with initial changes of +/-5 Hz and final changes on the magnitude of +/- 0.1 Hz. Participants performed the task until the Flicker-Fusion system had identified the participant’s CFF.

#### Questionnaires

Participants were asked to complete two questionnaires, the Comprehensive Autistic Trait Inventory (CATI; English et al., 2021) and the Prodromal Questionnaire-Brief (PQ-B; Loewy et al., 2005), so that we could assess the extent to which autistic and prodromal schizotypy traits in a typical college sample relate to variations in IAF and duration perception. Cognitive differences can vary widely in the Autism Spectrum Disorder (ASD) population and may be linked to variations in IAF (Dickinson et al., 2018). Research indicates that some individuals with ASD have atypical sensory processing, characterized by improved perception of local features, or detail, and impaired perception of global structure, or contextual information (Chung & Son, 2020; Dakin & Frith, 2005). However, there is conflicting evidence around whether individuals with ASD have typical or enhanced processing of time intervals (Poole et al., 2022; Wallace & Happé, 2008). Individuals with schizophrenia and schizotypal disorders, on the other hand, tend to have impairments in visual perception and visual working memory (Tek et al., 2002), which are thought to underlie difficulties in temporal processing and time perception (Roy et al., 2012), and have lower IAF on average (Ramsay et al., 2021; Sponheim et al., 2023; Trajkovic et al., 2021). Particularly, a meta-analysis suggests that individuals with schizophrenia are less accurate at discriminating durations and have the tendency to overestimate durations (Thoenes & Oberfeld, 2017). Thus, we wanted to explore varying amounts of autistic and schizotypal traits in our college sample related to variations in temporal perception.

To measure autistic traits in our college sample, we used the CATI (English et al., 2021), a 42-item inventory that asks participants to rate how much they agree with statements associated with traits typically seen in the ASD population. Responses are given via a 5-point scale ranging from “Definitely Disagree” to “Definitely Agree”. The items come from one of six main categories of traits (Social Interactions, Communication, Social Camouflage, Cognitive Rigidity, Repetitive Behavior, and Sensory Sensitivity) and include statements like, “Metaphors or ‘figures of speech’ often confuse me,” and “I feel discomfort when prevented from completing a particular routine.” Answers for each of the items were totaled for each participant, providing a score between 42-210.

The PQ-B (Loewy et al., 2005) was used to measure schizotypal or prodromal traits in the neurotypical population, an inventory consisting of 21 items asking about thoughts, feelings, and experiences within the past month. Items were responded to with a “yes” or “no” and included statements such as, “Do familiar surroundings sometimes seem strange, confusing, threatening or unreal to you?” and “Have you felt that you are not in control of your own ideas or thoughts?”. If the participant responded “yes”, a follow-up distress scale item asked whether the experience caused the participant to feel “frightened, concerned, or it causes problems” on a 5-point scale ranging from “Strongly Disagree” to “Strongly Agree”. Participant scores were computed in two ways: 1) by totaling the number of “yes” responses to get a score between 0-21, and 2) by totaling the values of the distress scale responses to get a score between 21-105. We were primarily interested in the first score which provided a numerical range of the prodromal experiences across individuals.

#### Procedure

Each day, participants signed a written consent form and filled out some basic questions about their state that day (e.g. tiredness, hours since they last ate, etc.) before completing additional tasks. On day one, participants answered demographic questions and completed the CATI and PQ-B questionnaires and the CFF task. Participants then completed at least two practice blocks of each duration task (more if their performance was low or participants reported having difficulty with the task) followed by one experimental block of each task where there was no feedback. On day two, participants quickly completed the CFF task before they were fitted with the EEG cap. They then completed one practice block and two experimental blocks of the duration estimation task, followed by one practice block and five experimental blocks of each duration discrimination task, with the static and dynamic task conditions counterbalanced. Finally, participants sat with their eyes closed for two minutes in order to collect resting state data at the end of the session. Participants were allowed to take breaks between blocks, as needed.

#### Behavioral Data Analysis

For the duration estimation task, we derived two standard measures of time estimation bias and precision by calculating each participant’s average estimates at each possible duration and their coefficient of variation (CV) in estimates at each possible duration, respectively. We calculated the CV for each participant and duration by dividing the standard deviation of participant estimates at each unique duration by the mean of their responses at each unique duration. The CV provided an unbiased measure of the variance in estimates, as it corrects for the larger variance inherent in estimates of longer durations. This measure can be interpreted as a precision measure (as variation in estimates decreases, precision increases). We also took the mean estimate and CV by collapsing across durations, so we had a single measure of estimation bias and precision for each participant.

Duration discrimination sensitivity was quantified from the discrimination tasks by fitting a logistic psychometric function (Palamedes toolbox version 1.10.4) to the proportion of times participants chose the comparison stimulus as “longer” for each of the possible comparison durations. This was done separately for each standard duration and each condition (static or dynamic). For all data fitting, the threshold parameter (reflecting the PSE) was bounded by the range of comparison values, the slope parameter of interest (β) was set to -10 to 100 to capture the expected positive slope, and the “lapse rate” parameters (lambda and gamma) values were fixed at 0.05 to denote the lower and upper bounds. Slopes were not normally distributed, so we applied a log10 transformation to each slope parameter. Thus, we derived a slope parameter as a measure of sensitivity for each participant and condition (100ms, 600ms, and 1200ms standards, static and dynamic conditions). Psychometric functions appeared to fit the duration discrimination data well for all participants and tasks except one participant who did not have a fit for the 1200ms dynamic condition. To confirm this, we computed R-squared values for the fit for each participant, standard duration, and stimulus type. The average goodness of fit across all participants, durations, and stimulus types was 0.86 (SD = 0.16; range: 0.78-0.93). When averaging across tasks, only one participant - the same participant whose data we were unable to fit for one condition - had an average goodness-of-fit below 0.50. Given that there was a correlation among slopes from the different tasks (see Figure 4B), we imputed the single missing datapoint by averaging the participant’s 1200ms static slope and 600ms dynamic slope.

#### EEG Data Collection

Continuous EEG was acquired from 63 active electrodes (BrainVision actiCHamp, iMotions A/S, Copenhagen, Denmark), with impedance kept below 20kΩ. Recordings were digitized at 1000 Hz, and FCz was used as the online reference. EEG was processed offline using custom scripts in MATLAB (version R2019b) and using EEGLAB toolbox (Delorme & Makeig, 2004).

EEG signals were first high-pass filtered at 0.1 Hz, downsampled to 500 Hz, and re-referenced using a median reference (to avoid noisy channels contaminating the reference at this stage of pre-processing). Trials were epoched based on the onset time of the stimulus (or the first stimulus in the discrimination task) to include 2s of prestimulus data and at least 800ms of post-second-stimulus data based on the longest possible stimulus duration(s) for each task. We also epoched the eyes-closed resting data into 105 1s-long epochs prior to manual inspection. All task data was then manually inspected to identify trials with muscular artifacts within a -500 prestimulus to +500 post-stimulus window, eye-blinks overlapping the stimulus presentations, and channels with excessive noise. After removing bad trials (estimation task: M = 14.62, SD = 19.83; discrimination task: M = 67.30, SD = 54.98) and interpolating noisy channels (estimation task: M = 2.71, SD = 1.76; discrimination task: M = 3.56, SD = 1.86) using spherical interpolation, an independent components analysis (ICA) was conducted, and ocular artifact components removed (M = 1.71, SD = 0.72). Eyes-closed data were manually inspected for noisy channels (M = 2.13, SD = 1.42) and epochs (M = 5.46, SD = 7.09) to be interpolated and rejected, respectively, but no ICA was run for this data. Finally, all data was average re-referenced and task data was baseline corrected using a 200ms prestimulus baseline window.

#### Individual Differences Analysis of IAF

A fast-Fourier transform (FFT) was used to identify peak alpha frequency for each individual from their eyes-closed resting data (Figure 2). First, each 1s epoch from the eyes-closed data was zero-padded, linearly detrended, and tapered using a Hamming window. After extracting power using the FFT, we averaged over an occipital electrode cluster (O1, O2, PO7, and PO8) with the highest group-level alpha power (see Figure 2). We then used the MATLAB function *findpeaks.m* to identify the frequency within the alpha-band range (7-14 Hz) that had the highest power for each participant, resulting in their IAF. If participants did not have a clear alpha peak (n=1), they were assigned a peak of 10 Hz. The average IAF of our sample was 10.04 Hz (SD = 0.96, range: 7.6 - 12.2 Hz). Note that we did not apply a logarithmic transform to the power spectrum, which is sometimes done to emphasize the 1/f slope of the power spectrum or help normalize power values across trials (Gyurkovics et al., 2021). Since we were primarily interested in obtaining a peak frequency per participant, the lack of log-transformation of the spectra actually helps to emphasize the alpha peaks (see Figure 2). However, the actual IAF values did not change as a result of this choice.

**Figure 2:**
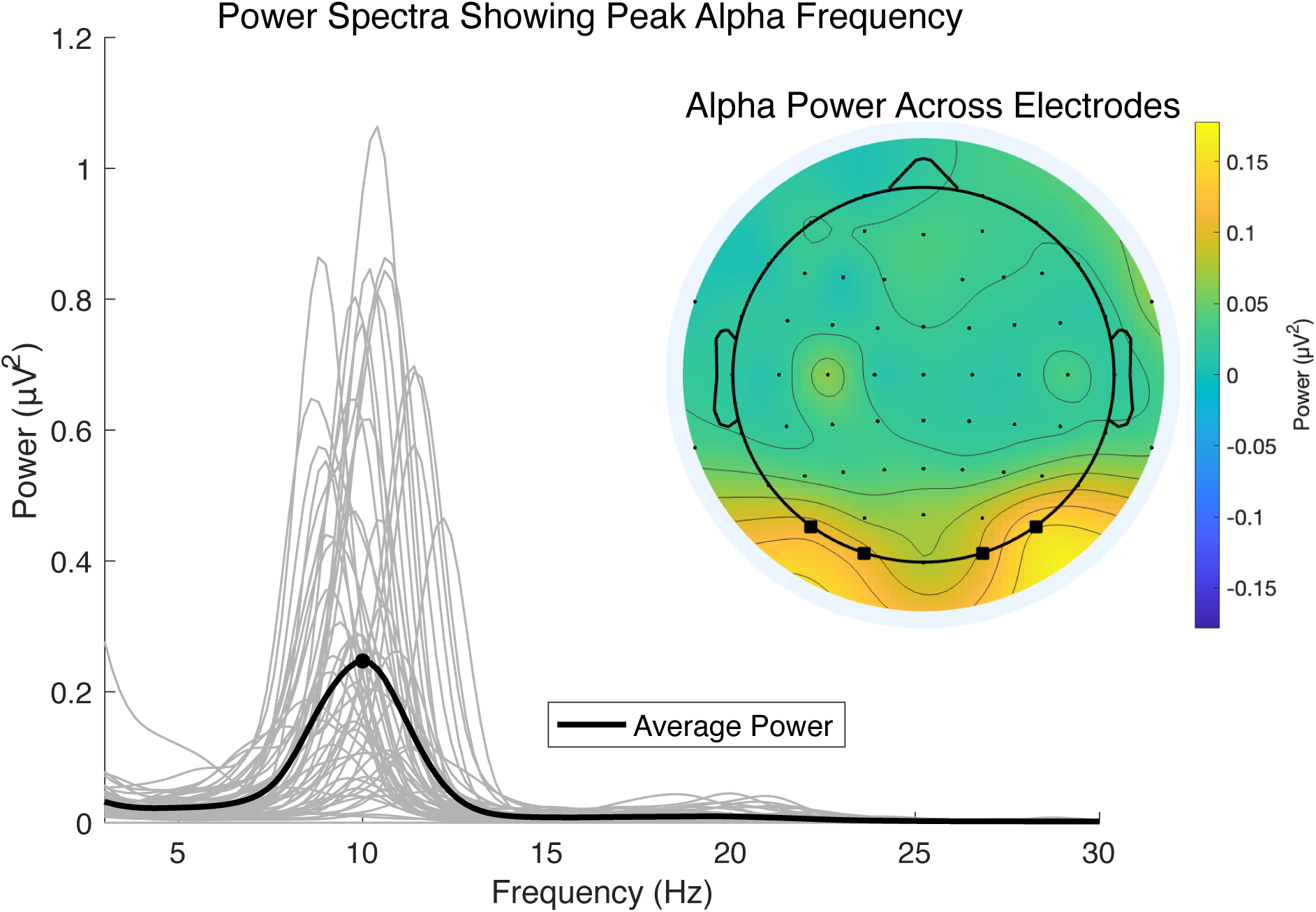
Individual and group power spectra from eyes-closed resting state data. The average power spectrum (black line) shows the average peak alpha frequency (black dot) is at 10 Hz. Gray lines are power spectra for individual subjects and emphasize the variability in peak frequencies. The topographical map shows the distribution of alpha power (7-14 Hz) across the scalp. Electrode locations (from left to right: PO7, O1, O2, PO8) with the highest group-level alpha power are highlighted with squares and were used in the FFT analysis to determine each individual’s peak alpha frequency.

We checked the relationship between eyes-closed IAF and trial-to-trial variations in task alpha frequency for each participant to determine how related the resting and task alpha frequencies were. This was done by subtracting alpha frequency estimates on single trials from the IAF defined at rest and then computing the mean and standard deviation of these differences, capturing the extent to which their instantaneous frequency differed from their resting-state IAF. Alpha frequencies were very similar across participants for all tasks. The estimation task had a mean frequency difference of 0.22 Hz (range: -0.76 - 0.36) and a mean standard deviation of 0.42 Hz (range: 0.27 - 0.53). Both static and dynamic tasks had, on average, a -0.10 Hz difference (range: -0.52 - 0.33 for both tasks) in trial frequency and participants’ IAF. The mean standard deviation for the static task was 0.29 Hz (range: 0.20 - 0.39) and for the dynamic task was 0.30 Hz (range: 0.22 - 0.38). These results suggest that while there may be trial-to-trial variability in alpha frequency, it is closely aligned with individual IAF and tends to vary around one’s IAF. Thus, we used eyes-closed IAF for all individual differences analyses since it provided the clearest and strongest measure of IAF.

Our main goal was to evaluate the relationship between IAF and duration perception across individuals while controlling for participant age (which could correlate with IAF, performance, or both) and while examining the dependence of any effect on the actual stimulus duration. Thus, beginning with the duration estimation data, we ran two mixed-effects regression models using stimulus duration, IAF (mean-centered), age (mean-centered), and the interaction of stimulus duration and IAF to predict mean estimates and the CV of estimates. To examine specific relationships more directly, we also used Spearman correlations to assess the relationship between IAF and behavior in the duration estimation task (namely the mean estimates and CV of estimates), after residualizing the behavioral measures for age.

For the duration discrimination task, we ran a mixed-effects regression model predicting duration discrimination slopes using the main effects of IAF (mean-centered), standard duration (100, 600,1200), stimulus type (static or dynamic; dummy-coded), participant age (mean-centered), and the interactions between IAF and standard duration, and IAF and stimulus type, as predictors. To examine specific relationships from the model we also used Spearman correlations to assess the relationship between IAF and the slopes for each of the conditions as well as the mean slopes across all conditions after correcting for age in each behavioral measure of performance using a linear model (both age-corrected and non-age corrected results can be found in Extended Data Table 4-1).

#### Trial-level Analysis of Alpha Frequency

We additionally explored the relevance of within-subject variation in alpha frequency using the “frequency sliding” approach developed in Cohen (2014), which measures the instantaneous alpha frequency at the single trial level (our implementation can be found in the MATLAB function *dotheslide.m* at https://samahalab.ucsc.edu/resources). We computed the instantaneous alpha frequency for each trial and electrode, averaged across -600 to -200ms prestimulus range (to minimize post-stimulus contamination) and around each participant’s IAF +/- 2 Hz. We reasoned that if IAF was linked to duration perception, this might also manifest in a link between trial-to-trial variation in alpha frequency and duration estimation or discrimination behavior. A higher or lower spontaneous prestimulus frequency may lead to a tendency to over- or under-estimate durations in the estimation task or lead to a change in sensitivity in the discrimination task. We also hypothesized that, in the discrimination task, a trial on which alpha frequency was higher prior to the comparison versus the standard stimulus might be more likely to result in a judgment that the comparison was longer, and thus result in a shift in the point of subjective equality (PSE) of the psychometric function.

We then performed a within-subjects analysis on the instantaneous alpha frequency extracted from each trial by employing a jackknife procedure on dependent measures from both tasks. To this end, we re-computed each measure for each jackknife sample (i.e., all trials excluding the current sample) and for each stimulus duration in the estimation task or each standard duration in the discrimination task. Jackknife sample measures were compared against measures computed with the full trial list, resulting in difference scores reflecting each trial’s contribution to each behavioral measure. We then used Spearman correlations between the alpha frequency measure of interest and the corresponding jackknife trial’s behavioral score. Specifically, the single-trial jackknife estimates of the CV of estimates and the mean estimate were correlated with the instantaneous frequency on each trial. The jackknife slope estimate for each trial of the discrimination task was correlated with the average instantaneous frequency prior to the standard and comparison stimuli. Finally, the jackknife PSE on each trial was correlated with the difference in instantaneous frequency, whereby we subtracted the comparison prestimulus frequency from the standard prestimulus frequency. Thus, we obtained a correlation value for each duration and participant and, for the discrimination task, each task condition. We then used a linear mixed-effects model to assess the role of duration and duration plus condition on the inverse hyperbolic tangents of correlations for the estimation measures and discrimination measures, respectively. The duration predictors for both models were minimum-centered so that the intercept represents the correlation between instantaneous frequency and behavior at the 300ms duration for the estimation task and the 100ms standard condition for the discrimination task. We fit random slopes for all models, and fit random intercepts for both duration and stimulus type in the discrimination measures models.

#### Exploratory Analysis

We were also curious about the extent to which CFF, CATI, and PQ-B scores related to IAF and duration perception sensitivity, as well as how the sensitivity measures across tasks related to one another. Given the non-normal distribution of CFF, CATI, and PQ-B scores, we computed separate Spearman correlations between IAF and each of these variables. We also used a Spearman correlation to explore the relationship between the CATI and PQ-B scores and participants’ mean estimates, CV of estimates, and duration discrimination slope values.

Additionally, we wanted to compare our main behavioral measures of interest across tasks to see how duration estimation related to duration discrimination, and to see how sensitivity measures related to bias measures. We computed a Spearman correlation between the mean slope of the duration discrimination task (taken across all conditions) and the CV of estimates as well as the mean slope of the duration discrimination task and the grand mean of estimates from the duration estimation task. Lastly, we checked whether the CV of estimates and mean of estimates from the estimation task and the mean slope and mean PSE from the duration discrimination task were correlated using a Spearman correlation.

## Results

### Duration Estimation: Individual Differences

We were primarily interested in looking at participants’ average estimates at each duration, as a measure of bias (Figure 3A, C, and E), and their variation around those estimates (CV), as a measure of precision (Figure 3B, D, and F). Mean duration reports (Figure 3E) were near veridical for medium durations (600-800ms) were underestimated at longer durations (900-1100ms), and overestimated at short duration (300-500ms), replicating a well-documented pattern in estimation and reproduction tasks thought to stem from a prior over durations centered on the mean of the stimulus distribution (Jazayeri & Shadlen, 2010). A mixed effect model of mean estimates (Figure 3A) demonstrated that stimulus duration was a significant predictor (β = 0.70, SE = 0.01, *t*(490) = 79.61, *p* < .001). As expected, as durations increased, so did the average duration estimates. No other predictors (age, IAF, or the interaction between IAF and duration) were significant in predicting mean estimation responses, suggesting IAF was not related to estimation bias. A mixed effect model of the CV of estimates (Figure 3B) found that CVs were also significantly predicted by stimulus duration (β = -0.001, SE < 0.001, *t*(490) = -11.91, *p* < .001) and IAF (β = -0.019, SE = 0.001, *t*(490) = -2.18, *p* = .03), but not by age or the interaction between IAF and duration. In other words, as durations increased, the CV of estimates decreased. Importantly, higher IAF was predictive of lower CV of estimates, indicating more precise time estimation responses. To help visualize this model we also computed correlations between IAF and age-corrected behavioral measures. We found that age-corrected mean estimates (Figure 3C) were not significantly related to IAF (*rho*(52) = .08, *p* = .57). However, the age-corrected CV of estimates (Figure 3D) was significantly negatively correlated with IAF (*rho*(52) = -.30, *p* = .03), such that faster IAF was associated with a lower CV, or greater precision in one’s time estimation. To visualize these relationships across stimulus durations, we also computed mean estimates and the CV of estimates for high and low IAF participants (median split), as shown in Figure 3E, F.

**Figure 3:**
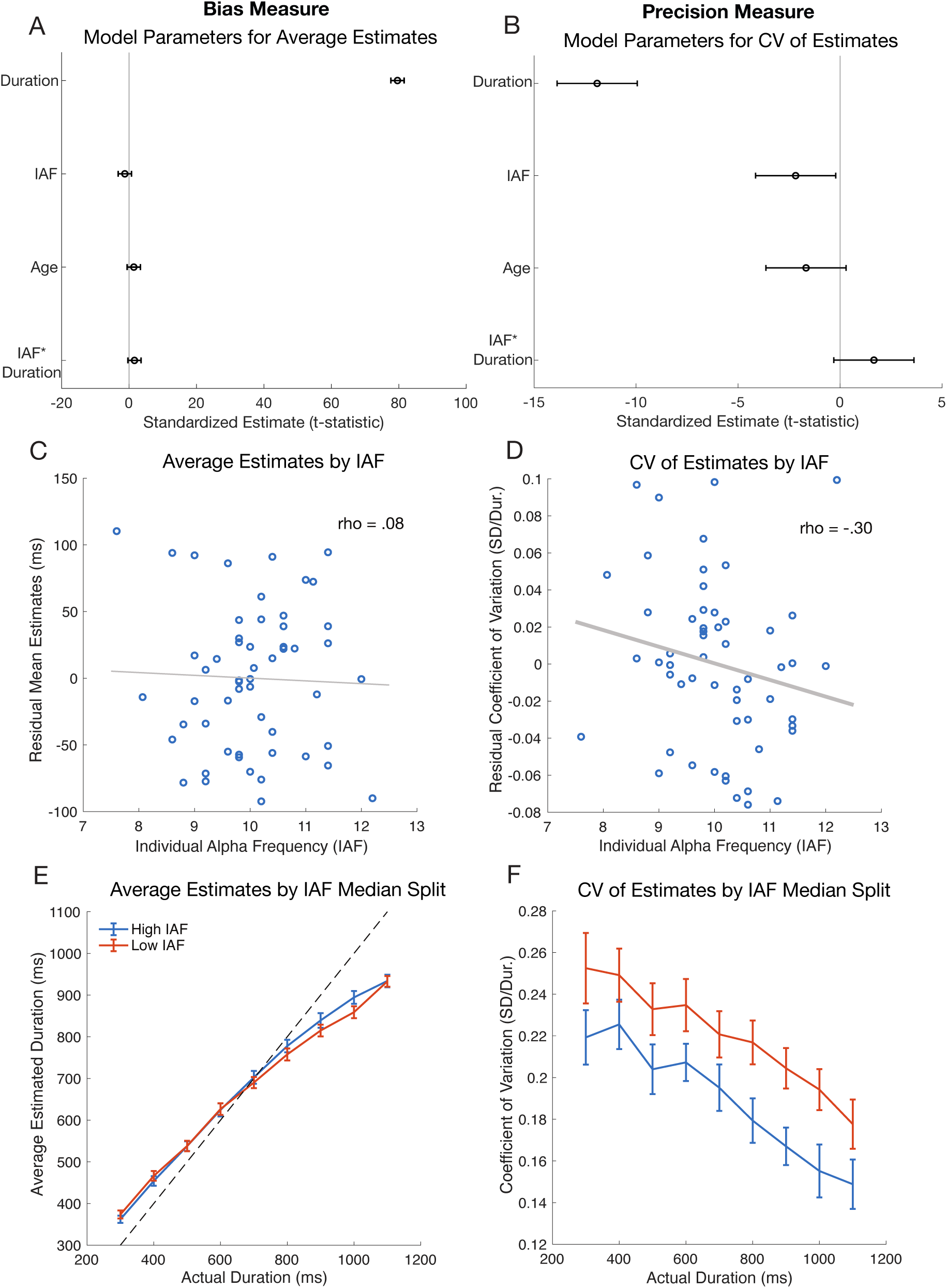
Duration estimation model output and measures, with average estimates (bias measures) plotted in the left column and the coefficient of variation (CV) of estimates (precision measures) plotted in the right column. (A-B) The effects (t-statistics and 95% CI) from linear mixed effects models that were used to predict average estimates and CV of estimates using Stimulus Duration, IAF, Age, and the interaction term IAF * Stimulus Duration. (C-D) Average estimates and CVs of estimates were averaged across all durations and corrected for age. Plots show the correlation between IAF and the residuals for mean estimates and CV of estimates, where each blue dot indicates an individual participant. Bolded fit lines indicate significance (p < .05). (E-F) Participants were split by the median IAF for visualization of the effect of IAF. Average estimates and CV of estimates are plotted for every actual duration that was presented in the duration estimation task. Participants with high IAF are shown in blue and participants with low IAF are shown in red (error bars are SEM).

#### Duration Discrimination: Individual Differences

Psychometric functions characterizing duration discrimination behavior were fit for each of the three standard durations (Figure 4A) and for low and high IAF individuals based on median split (Figure 4B). Performance in the discrimination task seemed to improve as durations became longer (Figure 4A), despite the comparison durations being proportionally matched to satisfy Weber’s law (Haigh et al., 2021), implying a breakdown of Weber’s law in duration discrimination at the shortest (100ms) duration. Despite this, we also found that individual differences in slopes across all standard durations were positively correlated with one another, ranging from *rho*(54) of .30-.79; see Fig. 4C).

**Figure 4:**
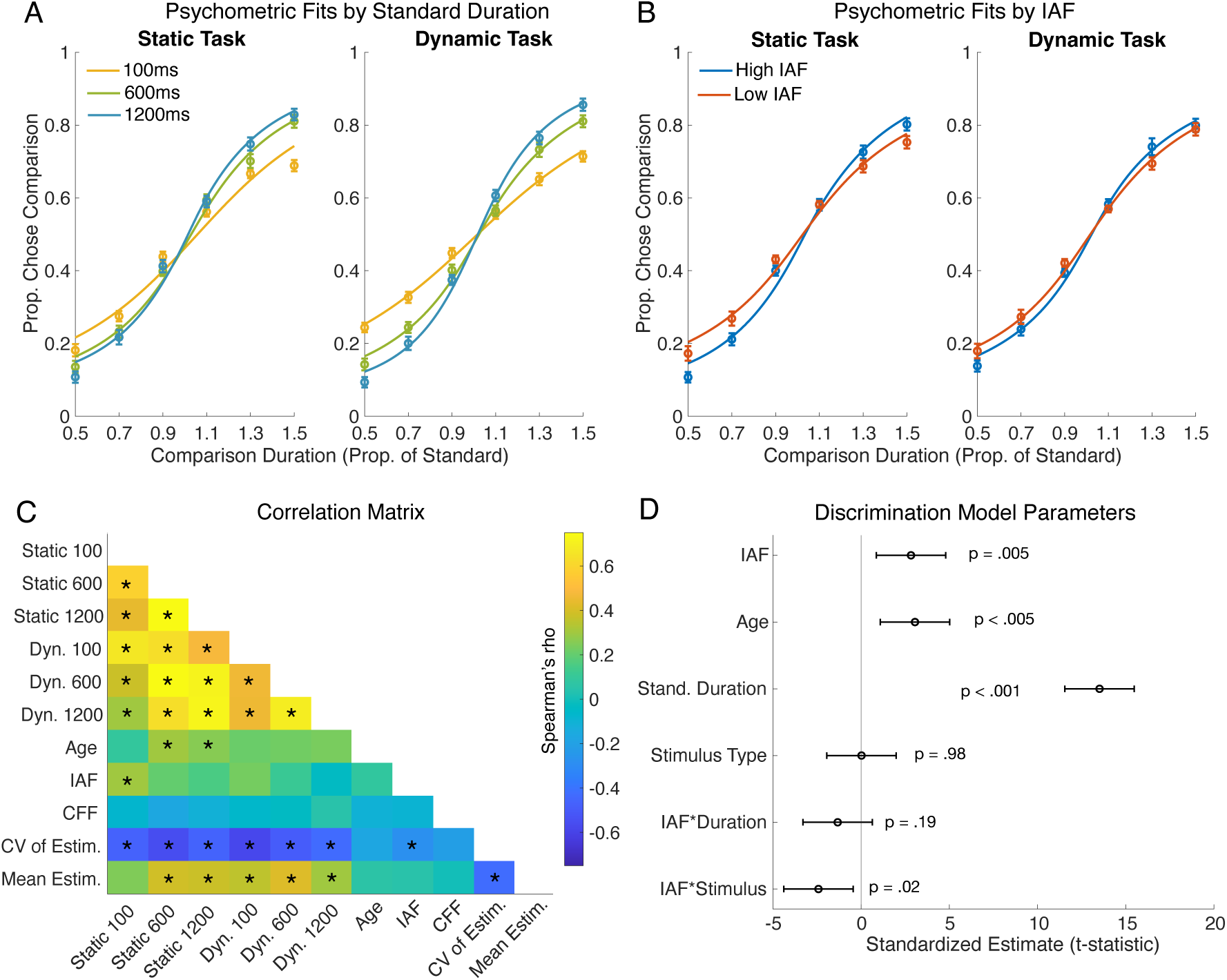
Duration discrimination results. (A) The proportion of times participants chose the comparison stimulus as being “longer” than the standard stimulus is plotted against the comparison duration (expressed in fractions of the standard). Psychometric function fits (lines) are shown alongside group mean data (circles with SEM error bars) for the 100ms standards (yellow line), 600ms standards (green line), and 1200ms standards (blue line), separated by stimulus type. (B) Psychometric functions for participants with high IAF (blue line) and low IAF (red line) according to a median split shown separately for the static and dynamic tasks. (C) A correlation matrix showing the Spearman correlation between duration discrimination slopes from each condition (from top to bottom, or left to right: Static 100ms, Static 600ms, Static 1200ms, Dynamic 100ms, Dynamic 600ms, Dynamic 1200ms), Age, IAF, CFF, the CV of Estimates (the sensitivity measure in the estimation task), and the Mean Estimate (the bias measure in the estimation task). Asterisks indicate significance (p < .05). Note that the effects in the correlation matrix are not corrected for age. (D) The effects (t-statistics and 95% CI) from a linear mixed effects model that was used to predict duration discrimination sensitivity (i.e., slopes) using IAF, Age, Standard Duration, Stimulus Type and the interaction terms IAF * Standard Duration and IAF * Stimulus Type in the model.

To estimate the joint effects of age, IAF, standard duration (100, 600, or 1200ms) and stimulus type (static versus dynamic) on psychometric slopes using a single model that incorporates all the conditions of the discrimination task, we estimated a linear mixed-effects model (see Methods and Fig. 4D). We found that participant performance could be predicted using IAF, age, standard duration, and stimulus type (static or dynamic) as predictor variables (R^2^ = .72). Specifically, age (β = 0.02, SE = 0.005, *t*(323) = 3.06, *p* = .002), IAF (β = 0.10, SE = 0.03, *t*(323) = 2.83, *p* = .005), and standard duration (β < 0.001, SE < 0.001, *t*(323) = 13.52, *p* < .001) were significant predictors of slope as a measure of task performance. While stimulus type (static or dynamic) was not a significant predictor of performance, it significantly interacted with IAF to predict performance (β = -0.05, SE = 0.02, *t*(323) = -2.42, *p* = .02), indicating that the relation between the duration discrimination sensitivity and IAF was stronger for static compared to dynamic stimuli (as can be seen in Figure 4B). There was no significant interaction between IAF and standard duration as a predictor variable, suggesting that the IAF relation to duration discrimination sensitivity was not strongly dependent on overall stimulus duration. Thus, when pooling together the discrimination sensitivity measures from all conditions into a single model, we found a clear positive association between IAF duration discrimination sensitivity that remained relatively constant across the three standard intervals tested (Fig. 4D). This effect indicates that individuals with higher IAF are more sensitive (i.e., have a steeper psychometric function slope) in discriminating between two different visual stimulus durations.

To help unpack and visualize the model results, we next explored the correlations between IAF and age-corrected duration discrimination sensitivity for different groupings of the discrimination task conditions. First, we evaluated whether IAF was related to duration discrimination sensitivity when averaging over all durations and stimulus types (Figure 5). We found that condition-averaged sensitivity (slope) was significantly positively correlated with IAF (*rho*(52) = .29, *p* = .03), such that individuals with faster alpha frequency exhibited more sensitive time perception (steeper psychometric function slopes, after controlling for age). Breaking down this relationship further, we found a significant correlation between IAF and the average slope for the static condition (*rho*(52) = .37, *p* < .01), but not for the dynamic condition (*rho*(52) = .13, *p* = .35), consistent with the interaction effect observed in the mixed-effects model (Fig. 4D). Finally, there was a significant correlation between IAF and sensitivity discriminating 100ms standards (*rho*(52) = .30, *p* = .02) and a moderate correlation between IAF and sensitivity discriminating 600ms standards (*rho*(52) = .26, *p* = .05), but no significant relationship was found for the 1200ms standards (*rho*(52) = .17, *p* = .22). Note, however, that these correlations are not statistically different from one another as reflected in the lack of a significant IAF-by-duration interaction in the mixed-model (Fig. 4D).

**Figure 5:**
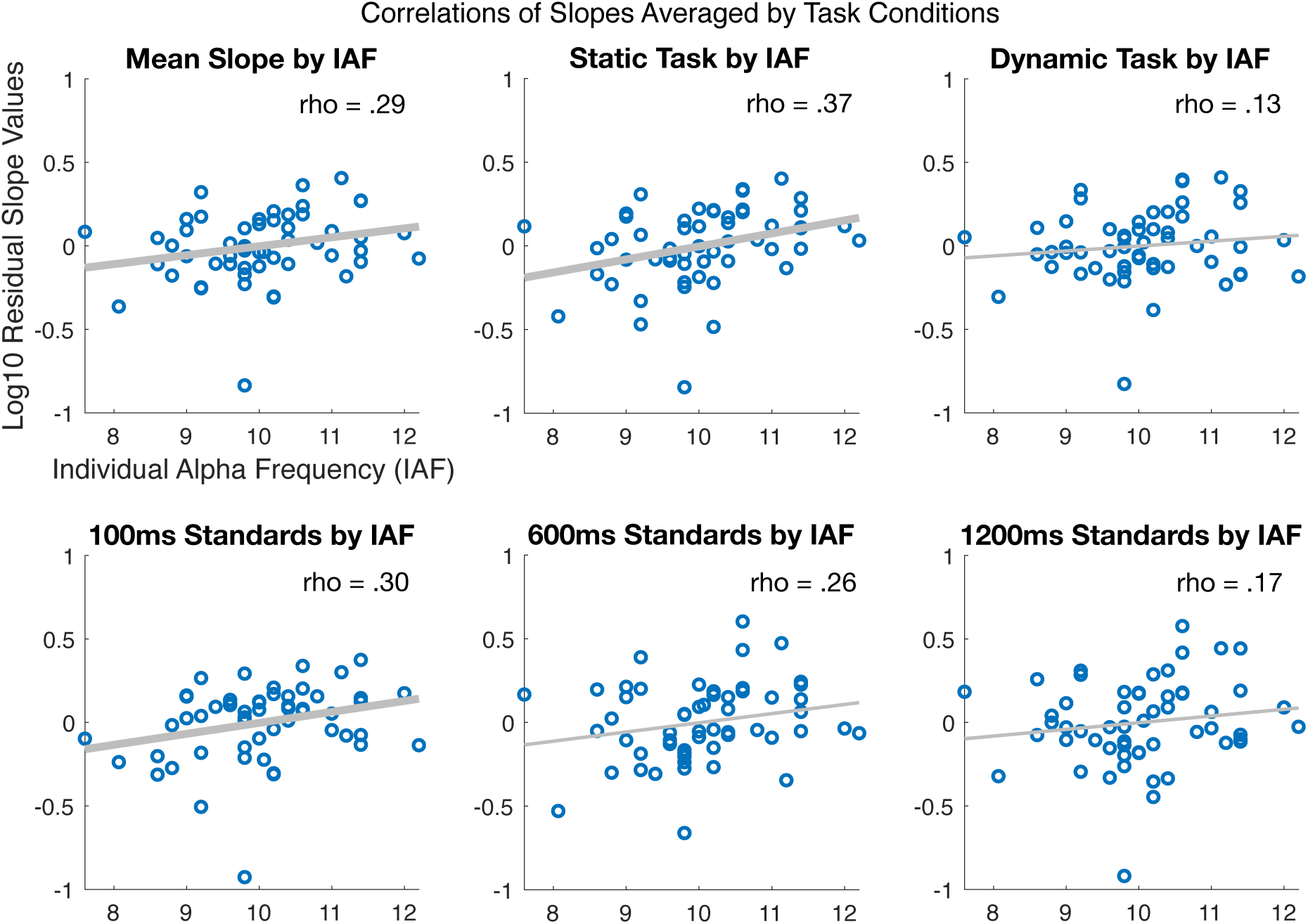
Correlations between IAF and (age-corrected) slopes for different duration discrimination task conditions. The average discrimination slope across all conditions is plotted in the top left, with stimulus types plotted in the middle and top rightmost plots (static condition and dynamic condition, respectively). Slopes from each standard duration (averaged over static and dynamic tasks) are plotted in the bottom row (from left to right: 100ms, 600ms, 1200ms) and). Each blue dot indicates an individual participant. Bolded fit lines indicate significance (p < .05).

#### Trial-level Measures

Finally, we were interested in how prestimulus instantaneous alpha frequency changes at the trial-level relate to bias and precision measures in duration estimation and discrimination (Figure 6A). For the estimation task, the mean correlation score across all subjects between single-trial prestimulus alpha frequency and (jackknifed) mean estimates was -0.01 (SD: 0.24) and was 0.02 (SD: 0.25) for the CV of estimates. The results of our mixed effects models indicate no significant correlations between pre-stimulus alpha frequency and mean estimates and no main effect of duration on these correlations. Additionally, there was no significant correlation between alpha frequency and the CV of estimates, however the model indicates a significant intercept term, reflecting the 300ms condition by which we minimum-centered the data (*t* = 2.37, *p* = 0.2).

**Figure 6.**
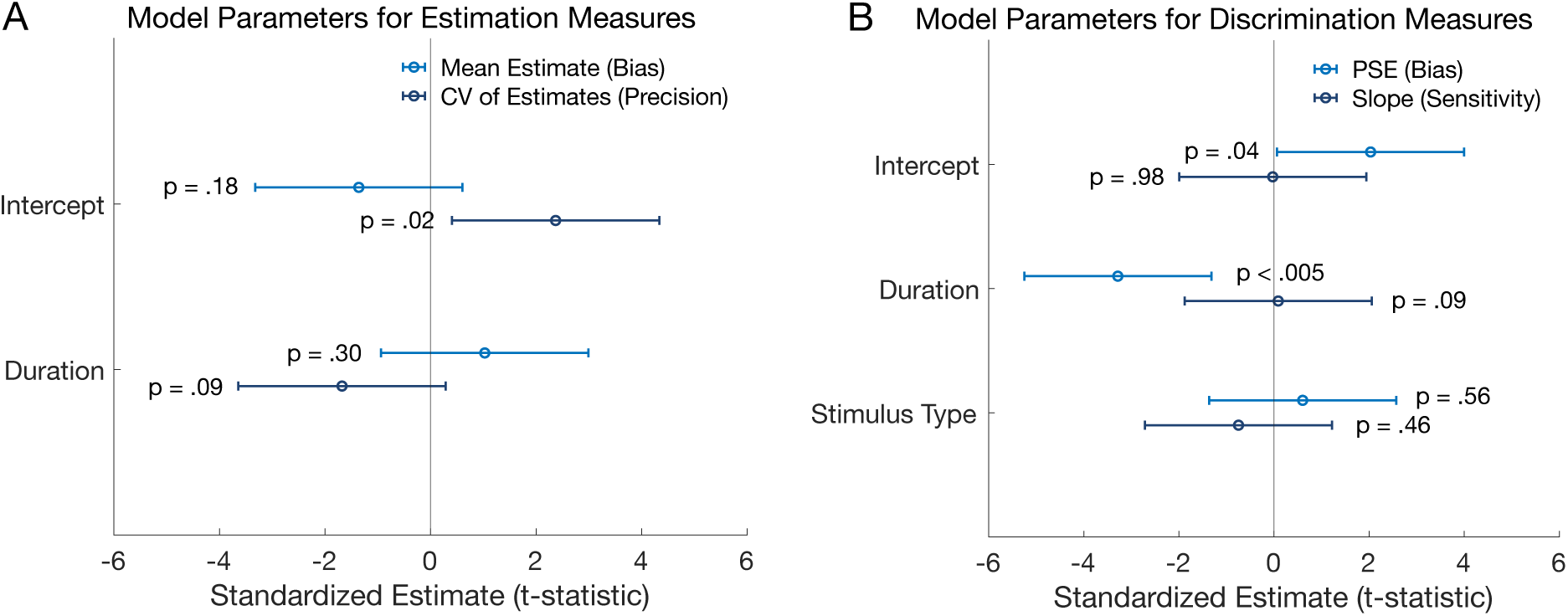
Results from linear mixed-effects models on the correlations between trial-level changes in instantaneous alpha frequency and measures from (A) the estimation task and (B) the discrimination task following a jackknife sampling procedure. Light blue lines (top rows of results) represent the bias measure correlations from each task: mean estimates (left) or the point of subjective equality (PSE; right) for choosing the standard as longer. Dark blue lines (bottom rows of results) indicate the sensitivity or precision measures: the CV of estimates (left) or the slope of psychometric fits (right). Intercept and duration are significant predictors of the relationship between instantaneous alpha frequency and PSE at the trial level.

We next used mixed models to evaluate the correlations obtained from the jackknife procedure between instantaneous prestimulus alpha frequency and discrimination behavior (Figure 6B). The mean correlations between the trial mean frequency and (jackknifed) slopes by condition were 0.0003 (SD: 0.08) for the static task and -.01 (SD: 0.09) for the dynamic task. The mean correlations between the difference in standard and comparison frequencies and (jackknifed) PSEs by condition were -0.005 (SD: 0.09) for the static task and -0.0005 (SD: 0.09) for the dynamic task. The model results indicated no main effect of correlation between trial frequency and slope. Additionally, neither duration nor stimulus type was a significant predictor of the frequency-slope correlations. The model of PSE and frequency differences, however, shows a significant intercept term, which represents the effect at the 100ms condition (*t* = 2.03, *p* = .04). Moreover, duration was a significant predictor of the correlation between PSE and frequency differences at the trial level (*t* = -3.29, *p* = .002), suggesting that the effect depended on duration and was specific to the 100ms condition. The direction of this effect implies that the PSE is shifted towards selecting the stimulus with a higher pre-stimulus alpha frequency, consistent with the idea that faster single-trial alpha produces inflated temporal estimates, but only for very brief stimuli.

#### Exploratory Measures

We were interested in evaluating whether IAF or duration perception performance related to questionnaire scores measuring Autistic-like traits and prodromal (schizotypal) experiences in each participant. Given our non-clinical population, we found generally low CATI and PB-Q scores (Extended Data Figure 4-1). The range of scores for the CATI was 73-187 (M = 116.16, SD = 26.50), and the range for the PB-Q was 0-18 (M = 4.89, SD = 4.45). No significant correlation was found between IAF and individual’s CATI scores or PQ-B scores. The CATI and PQ-B scores also did not significantly relate to any of the main performance measures of interest (mean estimates, CV of estimates, and duration discrimination slopes). Given the established relationship between CFF and IAF in clinical populations, we also compared CFF to various measures. First, we explored potential correlations between IAF with the CFF averaged across both testing days as well as the CFF from only the second day - the same testing session where we measured IAF. Neither CFF score was significantly correlated with IAF (average CFF: *rho*(53) = -.09, *p* = .51; day 2 CFF: *rho*(53) = -.15, *p* = .26). We also evaluated whether the CFF from the first day, the testing session where participants completed the trait questionnaires, was correlated with either questionnaire score (Extended Data Figure 4-1). We found a moderately weak, non-significant, correlation between CFF and CATI scores (*rho*(53) = .24, *p* = .08) and PQ-B scores (*rho*(53) = .23, *p* = .09).

Finally, we examined whether performance on the different discrimination tasks (estimation versus discrimination) was related across individuals. Since the CV derived from duration estimates and the slope measure derived from 2IFC discrimination are both sensitivity measures we would expect a relationship. Indeed, we found that the CV (averaged across durations) and the average slopes of the psychometric fits were significantly negatively correlated across participants (*rho*(53) = -.56, *p* < .001; Figure 4C), indicating that an individual with a steeper discrimination slope has lower estimation variability, as expected. Interestingly, there were also relationships with the bias measure derived from the estimation task such that participant’s average overall estimates correlated significantly with the average slopes of the psychometric fits (*rho*(53) = .41, *p* < .01).

## Discussion

This study measured individual differences in sensitivity and bias in duration perception using discrimination and estimation tasks across a range of peri-second durations. Our key finding is that IAF shows a medium-sized correlation with time perception sensitivity as measured by both estimation and discrimination tasks, and did not predict bias in duration estimation at the subject level. Importantly, our two sensitivity measures were significantly related: less variance in estimates was associated with a steeper psychometric function slope, highlighting a relationship between duration perception tasks that has not yet been explored. These effects hold when controlling for participant age, a confound that may have influenced conclusions in prior research. Our duration discrimination findings are consistent with, and expand upon, the vast literature supporting the role of IAF in sensitivity discriminating quick successive visual and multisensory stimuli (Cecere et al., 2015; Cooke et al., 2019; Di Gregorio et al., 2022; Migliorati et al., 2020; Noguchi, 2022; Samaha & Postle, 2015; Venskus & Hughes, 2021), and our duration estimation findings provide novel evidence that IAF also plays a role in the overall precision of estimating the duration of visual events.

We also show that IAF predicts time perception sensitivity approximately equally across a range of mostly sub-second durations, both in estimation and discrimination tasks. The shorter stimulus durations we tested were within what is traditionally considered a more “perceptual” duration judgment. The fact that our strongest correlation was found with 100ms standard, where Weber’s law was observed to break down, reinforces the idea that the alpha frequency supports “short” duration discrimination which likely relies even more on sensory mechanisms such as those involved in temporal resolution and temporal binding of stimuli. That said, we still found effects of IAF on sensitivity at our longer durations, which is supported by prior findings (Mokhtarinejad et al., 2024).

The role of IAF in duration discrimination was, however, specific to the static stimulus condition, as supported by a significant interaction between IAF and stimulus type in our mixed effects model. In our dynamic stimulus condition, we randomly modulated the luminance of the stimulus according to a white noise sequence, which has previously been shown to induce a strong alpha “echo” in the luminance-by-EEG cross correlation (VanRullen & Macdonald, 2012). We anticipated that this may enhance any link between IAF and duration discrimination although this was speculative, and we did not observe this effect. Instead, it’s possible these stimuli were more cognitively demanding to observe and duration perception in this task may have recruited more cognitive mechanisms. Alternatively, participants may have used different strategies for the dynamic task conditions such as trying to count notable luminance changes. Due to the nature of our blocked task conditions, we were unable to evaluate potential biases towards temporal perception of one stimulus type over another. This could be a compelling area of future research. Overall, it seems that the role of IAF in time perception may depend on the type of stimulus being judged.

We performed jackknife computations for trial-level analyses to explore whether spontaneous changes in alpha frequency relate to duration perception. We found no relationship between instantaneous prestimulus frequency bias (mean estimates) or precision (CV of estimates) in the estimation task, nor between the average prestimulus frequency and slope in the discrimination task. However, we did find a significant correlation between the difference in prestimulus frequency of the standard and comparison stimuli and PSE, as evidenced by a main effect of duration in our model, reflecting bias in the discrimination task. Specifically, as alpha frequency increased prior to the standard stimulus on the 100ms standard trials, participants were more likely to choose shorter comparison stimuli as equal in duration to the standard, indicating that faster trial-level alpha frequency makes short stimuli appear even shorter. Trial-to-trial variation in alpha frequency seems less clearly related to duration perception than IAF, given that the discrimination task duration effect did not generalize to the estimation task.

Some theories of time perception may explain individual trait-like alpha frequency relates to time perception sensitivity whereas spontaneous trial-by-trial fluctuations relate to discrimination bias (at short durations). The internal clock theory suggests that alpha provides a base temporal unit and that individuals adjust their temporal behaviors to their specific frequency (Karmarkar & Buonomano, 2007; Surwillo, 1966; Treisman, 1963; Treisman et al., 1990). Thus, individuals with faster alpha may get more visual information per unit of time, boosting temporal sensitivity, yet, through a lifetime of experience with a relative stable IAF, they are able to calibrate to reduce any tendency to under-or over-estimate time. However, when alpha fluctuates rapidly from trial-to-trial, this learned calibration may no longer hold and subjects may show subtle biases towards perceiving stimuli as longer if they occur during faster alpha states. However, single-trial instantaneous frequency changes are small and noisy, as reflected in our relatively low correlation values and the lack of a trial-level bias effect in the estimation data. Work by Kosciessa et al., (2020) has shown that an additional source of noise when estimating trial-level characteristics of oscillatory activity is inter-individual differences in signal-to-noise ratios, which we did not consider here. Here, we had hoped to reduce such noise with our jackknife analysis approach. However, the limited effects when relying on spontaneous variation in alpha frequency indicate the need for causal manipulations of alpha frequency at the within-subject level to test for a clearer relationship.

Our results are somewhat in conflict with recent work where IAF related to bias in temporal perceptions (Mioni et al., 2020), yet not sensitivity (Milton & Pleydell-Pearce, 2016; Mioni et al., 2020). However, the standard stimulus in these tasks was either learned before viewing comparisons, or presented as the first stimulus. Perceptual training research (Navarra et al., 2005; Powers et al., 2009; Stevenson et al., 2013) and research on the temporal order effect (Grondin, 2010) suggest that these design considerations have important implications for participant performance. Additionally, the bias effect emerged from tACS, which aligns our trial-level PSE bias effect and the idea that we may be calibrated to our IAF, and not to other alpha frequencies. Interestingly, research using a similar discrimination design did indeed demonstrate a role of alpha frequency in sensitivity, while also showing a role of alpha power in accuracy (Mokhtarinejad et al., 2024). Given conflicting findings, we note that our sample size was considerably larger than those prior papers and we found consistent results across two distinct time perception tasks and across different stimulus durations. However, it could be valuable to evaluate performance on these temporal perception tasks in relation to other alpha characteristics which might be underlying perceptual biases, such as phase, alpha bursts, or changes in alpha power, as seen in the prior literature (Azizi et al., 2023; Cahoon, 1969; Legg, 1968; Milton & Pleydell-Pearce, 2016; Mokhtarinejad et al., 2024; Werboff, 1962).

The final analyses examining IAF’s relationship to CFF, CATI, and PQ-B scores, and the CATI and PQ-B scores to task performance were somewhat exploratory in nature, and we failed to find any significant relationships across the measures. While IAF correlates with CFF in patients with hepatic encephalopathy (Baumgarten et al., 2018; May et al., 2014), it is possible that CFF and IAF are uniquely correlated in these clinical populations, perhaps due to other underlying mechanisms driving these patients to have lower IAF on average compared to healthy controls (Butz et al., 2013; Götz et al., 2013; May et al., 2014). The lack of correlation between IAF and CFF in our study fits with a recent paper showing IAF correlated with another temporal resolution measure (the two-flash fusion paradigm) but not with CFF, suggesting these measures tap different aspects of temporal resolution (Haarlem et al., 2024). Finally, despite IAF and duration perception variations in populations with ASD (Dickinson et al., 2018; Poole et al., 2022; Wallace & Happé, 2008) and schizotypal disorders (Ramsay et al., 2021; Roy et al., 2012; Sponheim et al., 2023; Tek et al., 2002; Thoenes & Oberfeld, 2017; Trajkovic et al., 2021), we found no significant correlations between IAF and the magnitude of ASD traits or prodromal traits, perhaps due to the non-normal distribution of trait scores in our population (see Extended Data Figure 4-1).

Overall, our results support the idea that rhythmic alpha sampling drives sensitivity in time perception, as faster IAF theoretically results in more frequent excitatory perceptual windows and updating of sensory information to be used for estimating duration. Further supporting this theory was the finding that the sensitivity measures were significantly negatively correlated across tasks: as discrimination sensitivity increased, the amount of variance in estimates decreased. While this correlation is not sufficient to conclude that these processes are supported by the same underlying mechanism, it is a necessary relationship to observe if IAF is indeed driving perceptual sensitivity across tasks. Finally, we note that many important future directions stem from this work, such as considering the role of IAF in filled versus empty intervals, as has often been done in behavioral duration perception studies (Buffardi, 1971; Hasuo et al., 2014; Wearden et al., 2007; Williams et al., 2019), and in non-visual or multimodal duration perception. Critically, though, causal methods are needed to establish the mechanistic role of alpha oscillations in time perception.

## Acknowledgments

This publication was supported in part by the National Institute on Drug Abuse, Award Number T32DA007250. The content is solely the responsibility of the authors and does not necessarily represent the official views of the National Institutes of Health.

## Extended Data

**Table 4-1:**
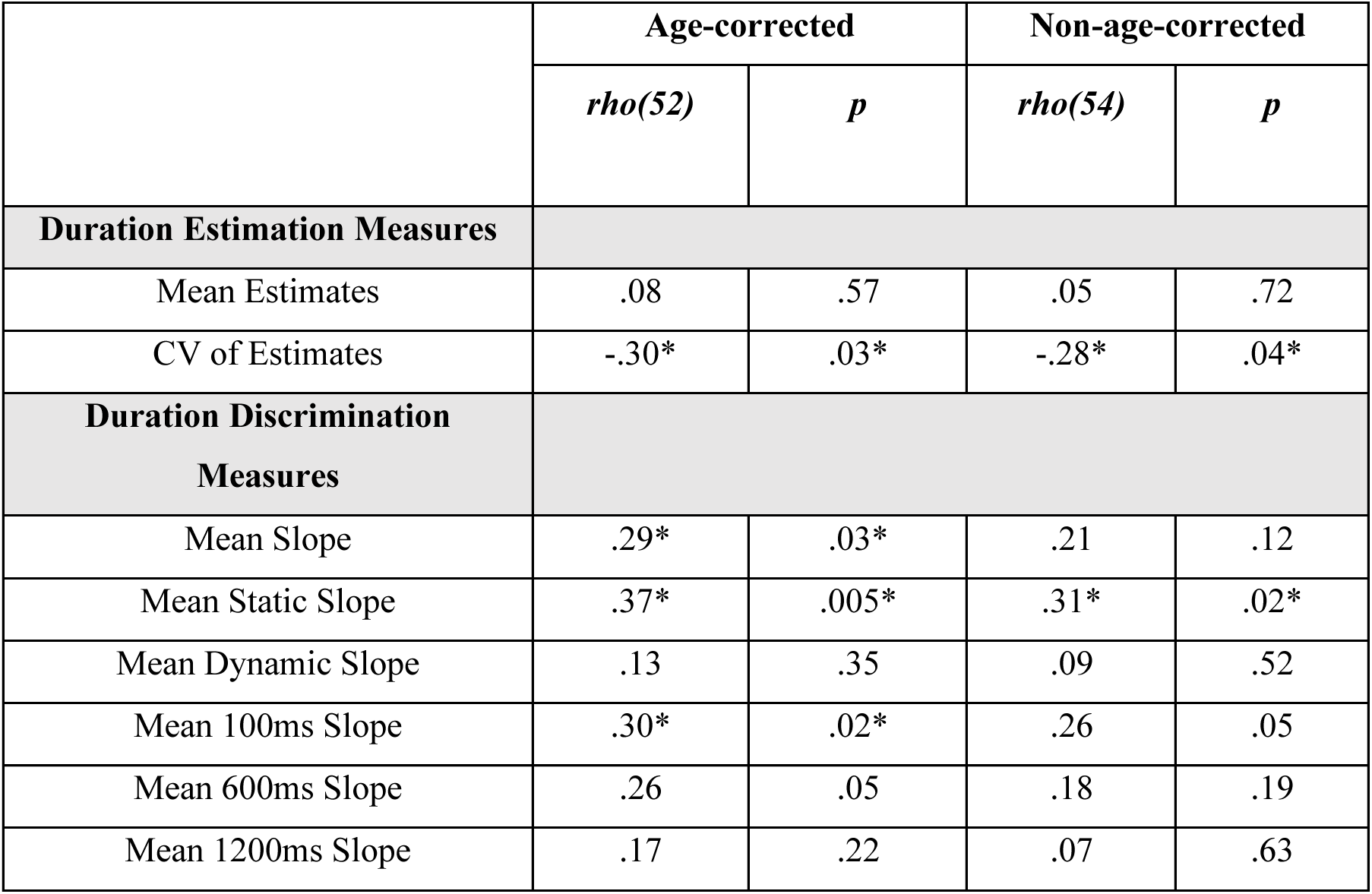
Age-corrected and non-age-corrected statistical results of duration estimation and duration discrimination analyses. Each measure was corrected for age using a linear model to account for the amount of variance explained in the performance measure by participant age. These correlations are shown in the first two columns of the table and displayed graphically in Figure 3C and D and Figure 5. Non-age-corrected correlations were also performed and are provided in the final two columns of the table, and are used in the correlation matrix of results shown in Figure 4C. Asterisks indicate significant correlations (p < .05).

**Figure 4-1.**
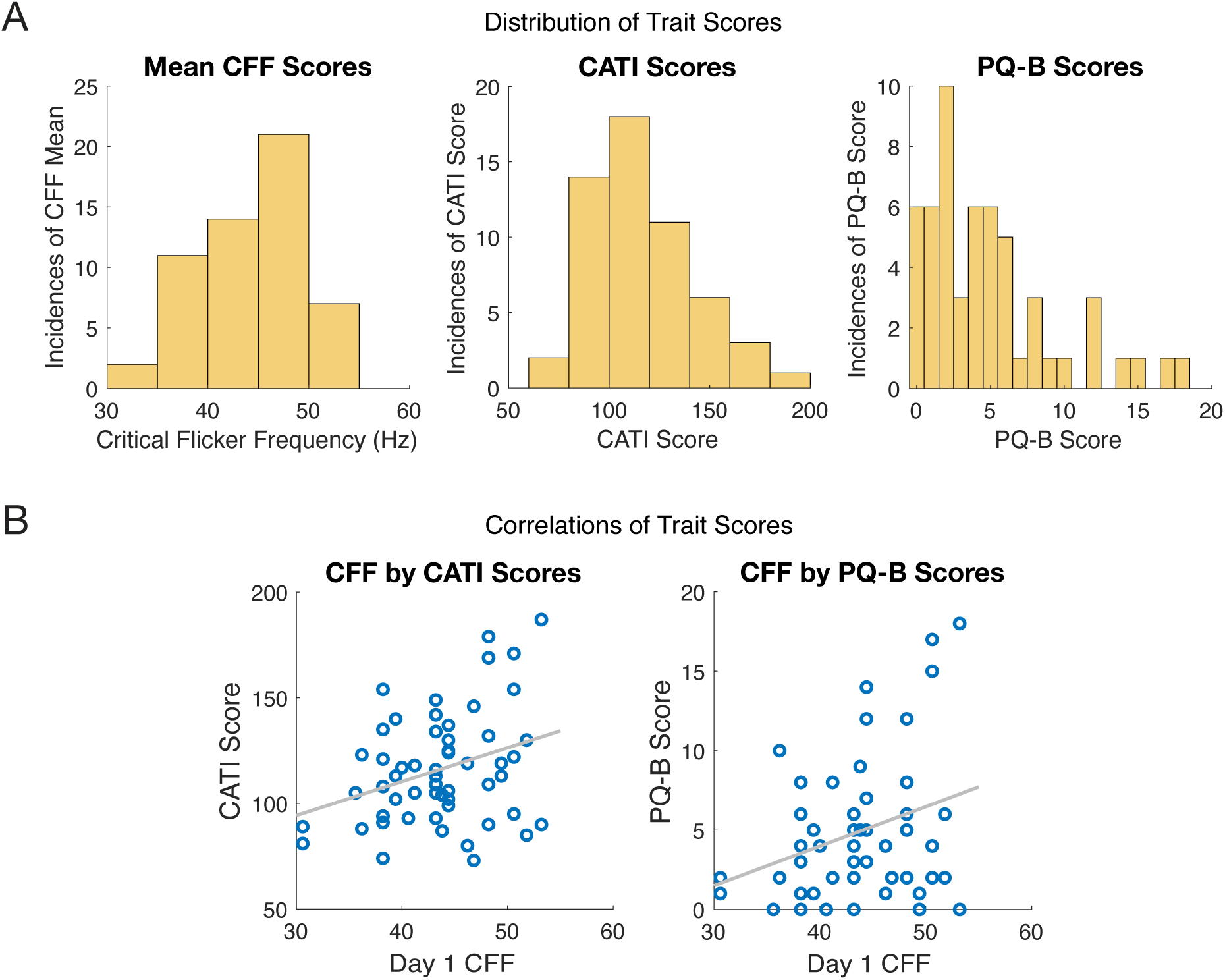
The distribution of CFF, CATI, and PQ-B scores across the sample, and correlations between Day 1 CFF and the CATI and PQ-B scores, which expand upon correlations from Figure 4C. (A) Histograms demonstrate the number of participants with each trait-like score that was measured in the experiment: critical flicker frequency (CFF), Comprehensive Autistic Trait Inventory (CATI) and the Prodromal Questionnaire (PQ-B). (B) A Spearman correlation was computed between the Day 1 CFF scores and the CATI scores (left) and PQ-B scores (right) as an exploratory measure of the relationship between traits that were expected to, but did not relate to, IAF. Neither correlation reached statistical significance.

